# RNAmountAlign: efficient software for local, global, semiglobal pairwise and multiple RNA sequence/structure alignment

**DOI:** 10.1101/389312

**Authors:** Amir H Bayegan, Peter Clote

## Abstract

Alignment of structural RNAs is an important problem with a wide range of applications. Since function is often determined by molecular structure, RNA alignment programs should take into account both sequence and base-pairing information for structural homology identification. A number of successful alignment programs are heuristic versions of Sankoff’s optimal algorithm. Most of them require *O*(*n*^4^) run time. This paper describes `C++` software, `RNAmountAlign`, for RNA sequence/structure alignment that runs in *O*(*n*^3^) time and *O*(*n*^2^) space; moreover, our software returns a *p*-value (transformable to expect value *E*) based on Karlin-Altschul statistics for local alignment, as well as parameter fitting for local and global alignment. Using incremental mountain height, a representation of structural information computable in cubic time, `RNAmountAlign` implements quadratic time pairwise local, global and global/semiglobal (query search) alignment using a weighted combination of sequence and structural similarity. `RNAmountAlign` is capable of performing progressive multiple alignment as well. Benchmarking of `RNAmountAlign` against `LocARNA, LARA, FOLDALIGN, DYNALIGN` and `STRAL` shows that `RNAmountAlign` has reasonably good accuracy and much faster run time supporting all alignment types.

**Availability:** `RNAmountAlign` is publicly available at http://bioinformatics.bc.edu/clotelab/RNAmountAlign.

## Introduction

A number of different metrics exist for comparison of RNA secondary structures, including base pair distance (BP), string edit distance (SE) [1], mountain distance (MD) [2], tree edit distance (TE) [3], coarse tree edit distance (HTE) [4], morphological distance [5] and a few other metrics. In what appears to be the most comprehensive published comparison of various secondary structure metrics [6], it was shown that all of these distance measures are highly correlated when computing distances between structures taken from the Boltzmann low-energy ensemble of secondary structures [7] for the same RNA sequence – so-called *intra-ensemble* correlation. In contrast, these distance measures have low correlation when computing distances between structures taken from Boltzmann ensembles of different RNA sequences of the same length – so-called *inter-ensemble* correlation. For instance, the intra-ensemble correlation between base pair distance (BP) and mountain distance (MD) is 0.822, while the corresponding inter-ensemble correlation drops to 0.210. Intra-ensemble correlation between string edit distance (SE) and the computationally more expensive tree edit distance (TE) is 0.975, while the corresponding intra-ensemble correlation drops to 0.590 – see Table 1.

**Table 1.** Correlation between various secondary structure metrics, as computed in [6]: base pair distance (BP), string edit distance (SE) [1], mountain distance (MD) [2], tree edit distance (TE) [3] and coarse tree edit distance (HTE) [4]. Lower triangular values indicate intra-ensemble correlations; upper triangular values indicate inter-ensemble correlations. Table values are taken from [6].

|  | BP | MD | SE | TE | HTE |
| --- | --- | --- | --- | --- | --- |
| BP |  | 0.210 | 0.134 | 0.133 | 0.230 |
| MD | 0.822 |  | 0.519 | 0.607 | 0.515 |
| SE | 0.960 | 0.853 |  | 0.590 | 0.310 |
| TE | 0.943 | 0.879 | 0.975 |  | 0.597 |
| HTE | 0.852 | 0.844 | 0.879 | 0.913 |  |

Due to poor inter-ensemble correlation of RNA secondary structure metrics, and the fact that most secondary structure pairwise alignment algorithms depend essentially on some form of base pair distance, string edit distance, or free energy of common secondary structure, we have developed the first RNA sequence/structure pairwise alignment algorithm that is based on (incremental ensemble) mountain distance. Our software, `RNAmountAlign`, uses this distance measure, since the Boltzmann ensemble of all secondary structures of a given RNA of length *n* can represented as a length *n* vector of real numbers, thus allowing an adaptation of fast sequence alignment methods. Depending on the command-line flag given, our software, `RNAmountAlign` can perform pairwise alignment, (Needleman-Wunsch global [8], Smith-Waterman local [9] or semiglobal [10] alignment) as well as progressive multiple alignment (global and local), computed using a guide tree as in `CLUSTAL` [11]. Expect values *E* for local alignments are computed using Karlin-Altschul extreme-value statistics [12,13], suitably modified to account for our new sequence/structure similarity measure. Additionally, `RNAmountAlign` can determine *p*-values (hence *E*-values) by parameter fitting for the normal (ND), extreme value (EVD) and gamma (GD) distributions.

We benchmark the performance of `RNAmountAlign` on pairwise and multiple global sequence/structure alignment of RNAs against the widely used programs `LARA, FOLDALIGN, DYNALIGN, LocARNA` and `STRAL. LARA` (Lagrangian relaxed structural alignment) [14] formulates the problem of RNA (multiple) sequence/structure alignment as a problem in integer linear programming (ILP), then computes optimal or near-optimal solutions to this problem. The software `FOLDALIGN` [15-17], and `DYNALIGN` [18] are different *O*(*n*^4^) approximate implementations of Sankoff’s *O*(*n*^6^) optimal RNA sequence/structure alignment algorithm. `FOLDALIGN` sets limits on the maximum length of the alignment as well as the maximum distance between subsequences being aligned in order to reduce the time complexity of the Sankoff algorithm. `DYNALIGN` [18] implements pairwise RNA secondary structural alignment by determining the common structure to both sequences that has lowest free energy, using a positive (destabilizing) energy heuristic for gaps introduced, in addition to setting bounds on the distance between subsequences being aligned. In particular, the only contribution from nucleotide information in `Dynalign` is from the nucleotide-dependent free energy parameters for base stacking, dangles, etc. `LocARNA` (local alignment of RNA) [19,20] is a heuristic implementation of `PMcomp` [21] which compares the base pairing probability matrices computed by McCaskill’s algorithm. Although the software is not maintained, `STRAL` [22] which is similar to our approach, uses up- and downstream base pairing probabilities as the structural information and combines them with sequence similarity in a weighted fashion.

`LARA, mLocARNA` (extension of `LocARNA`), `FOLDALIGNM` [16,23] (extension of `FOLDALIGN`), `Multilign` [24,25] (extension of `DYNALIGN`) and `STRAL` support multiple alignment. `LARA` computes all pairwise sequence alignments and subsequently uses the `T-Coffee` package [26] to construct multiple alignments. Both `FOLDALIGNM` and `mLocARNA` implement progressive alignment of consensus base pairing probability matrices using a guide tree similar to the approach of `PMmulti` [21]. For a set of given sequences, `Multilign` uses `DYNALIGN` to compute the pairwise alignment of a single fixed index sequence to each other sequence in the set, and computes a consensus structure. In each pairwise alignment, only the index sequence base pairs found in previous computations are used. More iterations in the same manner with the same index sequence are then used to improve the structure prediction of other sequences. The number of pairwise alignments in `Multilign` is linear with respect to the number of sequences. `STRAL` performs multiple alignment in a fashion similar to `CLASTALW` [27]. Table 2 provides an overview of various features, to the best of our knowledge, supported by the software benchmarked in this paper.

**Table 2.** Overview of features in software used in benchmarking tests, where ✓ [resp. —] indicates the presence [resp. absence] of said feature, to the best of our knowledge. Average F1 [resp. SPS] scores for the pairwise [resp. multiple] global alignment are given, computed as explained in the text.

| Software | Local | Global | Semiglobal | E-value | F1(Pairwise) | SPS(Multiple) |
| --- | --- | --- | --- | --- | --- | --- |
| RNAmountAlign | ✓ | ✓ | ✓ | ✓ | 0.84 | 0.84 |
| LocARNA | ✓ | ✓ | — | — | 0.81 | 0.84 |
| LARA | — | ✓ | — | — | 0.84 | 0.85 |
| FOLDALIGN | ✓ | ✓ | — | ✓ | 0.80 | 0.77 |
| DYNALIGN | — | ✓ | — | — | 0.68 | 0.67 |
| STRAL | — | ✓ | — | — | 0.82 | - |

`RNAmountAlign` can perform semiglobal alignments in addition to global and local alignments. As in the RNA tertiary structural alignment software `DIAL` [28], semiglobal alignment allows the user to perform a query search, where the query is entirely matched to a local portion of the target. Quadratic time alignment using affine gap cost is implemented in `RNAmountAlign` using the Gotoh method [29] with the following pseudocode, shown for the case of semiglobal alignment. Let *g*(*k*) denote an affine cost for size *k* gap, defined by *g*(0) = 0 and *g*(*k*) = *g_i_* + (*k* – 1) · *g_e_* for positive gap initiation [resp. extension] costs *g_i_* [resp. *g_e_*]. For query **a** = *a*_1_,…, *a_n_* and target **b** = *b*_1_,…, *b_m_*, define (*n* + 1) × (*m* + 1) matrices *M, P, Q* as follows: *M*_*i*,0_ = *g*(*i*) for all 1 ≤ *i* ≤ *n*, *M*_0,*j*_ = 0 for all 1 ≤ *j* ≤ *m*, while for positive *i, j* we have *M_i,j_* = max (*M*_*i*–1,*j*–1_ + sim (*a_i_*, *b_j_*), *P_i,j_*, *Q_i,j_*). For 1 ≤ *i* ≤ *n*, 1 ≤ *j* ≤ *m*, let *P*_0,*j*_ = 0 and *P_i,j_* = max (*M*_*i*–1,*j*_ + *g_i_*, *P*_*i*–1,*j*_ + *g_e_*), and define *Q*_*i*,0_ = 0 and *Q_i,j_* = max (*M*_*i,j*–1_ + *g_i_*, *Q*_*i,j*–1_ + *g_e_*, 0). Determine the maximum semiglobal alignment score in row *n*, then perform backtracking to obtain an optimal semiglobal (or query search) alignment.

In this paper we provide a very fast, comprehensive software package capable of pairwise/multiple local/global/semiglobal alignment with *p*-values and *E*-values for statistical significance. Moreover, due to its speed and relatively good accuracy, the software can be used for whole-genome searches for homologues of a given orphan RNA as query. This is in contrast to `Infernal` [30], which requires a multiple alignment to construct a covariance model for whole-genome searches.

## Materials and methods

### Incremental ensemble expected mountain height

Introduced in [31], the *mountain height*^1^ *h_s_*(*k*) of secondary structure *s* at position *k* is defined as the number of base pairs in *s* that lie between an external loop and *k*, formally given by

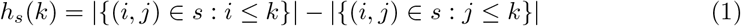

The *ensemble mountain height* 〈*h*(*k*)〉 [32] for RNA sequence **a** = *a*_1_,…,*a_n_* at position *k* is defined as the average mountain height, where the average is taken over the Boltzmann ensemble of all low-energy structures *s* of sequence **a**. If base pairing probabilities *p_i,j_* have been computed, then it follows that

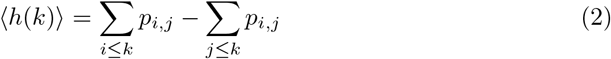

and hence the *incremental ensemble mountain height*, which for values 1 < *k* ≤ *n* is defined by *m*_a_(*k*) = 〈*h*(*k*)〉 – 〈*h*(*k* – 1)〉 can be readily computed by

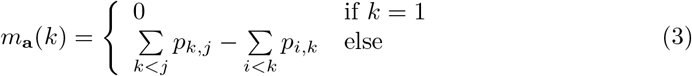

It is clear that —1 ≤ *m*_a_(*k*) ≤ 1, and that both ensemble mountain height and incremental ensemble mountain height can be computed in time that is quadratic in sequence length *n*, provided that base pairing probabilities *p_i,j_* have been computed. Except for the cubic time taken by a function call of `RNAfold` from Vienna RNA package [4], the software `RNAmountAlign` has quadratic time and space requirements. Figure 1 depicts a global alignment of two transfer RNAs, computed by `RNAmountAlign`, shown as superimposed ensemble mountain height displays with gaps.

**Fig 1.**
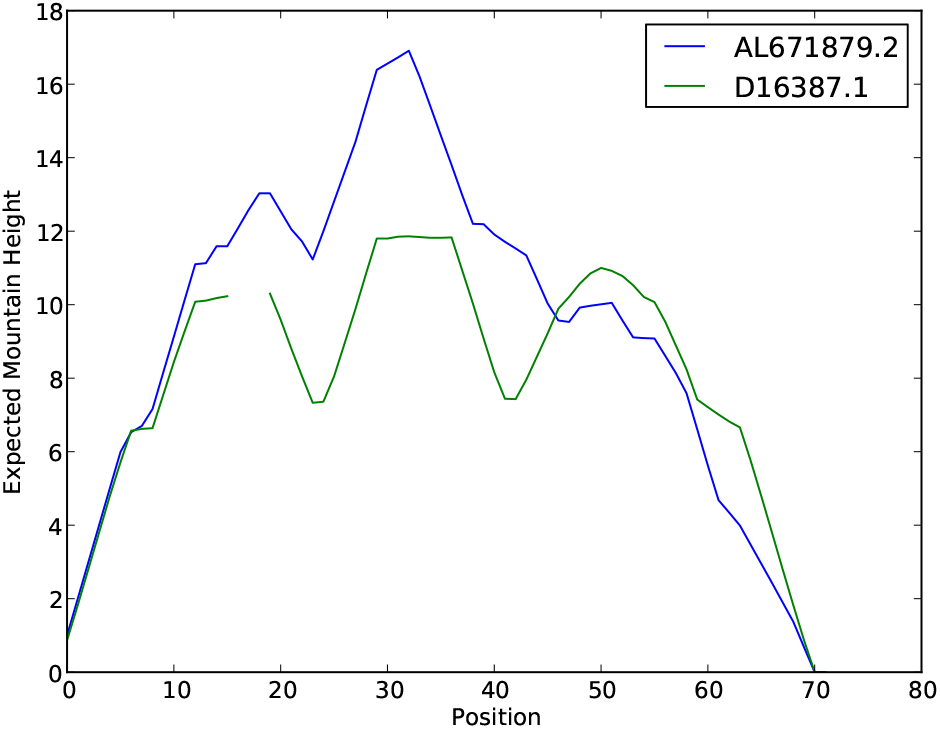
Ensemble mountain heights of 72 nt tRNA AL671879.2 and 69 nt tRNA D16387.1, aligned together by `RNAmountAlign`. Since the `BRAliBase 2.1 K2` reference (pairwise) alignment [33] has only 28% sequence identity, structural similarity parameter *γ* was set to 1 in our software `RNAmountAlign`, which returned the correct alignment. See Methods section for explanation of *γ* and the algorithm used by `RNAmountAlign`.

### Transforming distance into similarity

In [34], Seller’s (distance-based) global pairwise alignment algorithm [35] was rigorously shown to be equivalent to Needleman and Wunsch’s (similarity-based) global pairwise alignment algorithm [8]. Recalling that Seller’s alignment distance is defined as the minimum, taken over all alignments of the sum of distances *d*(*x, y*) between aligned nucleotides *x, y* plus the sum of (positive) weights *w*(*k*) for size *k* gaps, while Needleman-Wunsch alignment similarity is defined as the maximum, taken over all alignments of the sum of similarities *s*(*x, y*) between aligned nucleotides *x, y* plus the sum of (negative) gap weights *g*(*k*) for size *k* gaps, Smith and Waterman [34] show that by defining

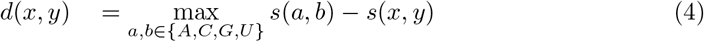

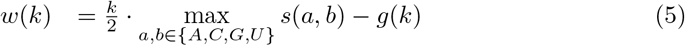

and by taking the minimum distance, rather than maximum similarity, the Needleman-Wunsch algorithm is transformed into Seller’s algorithm. Though formulated here for RNA nucleotides, equivalence holds over arbitrary alphabets and similarity measures (e.g. BLOSUM62).

For *x, y* ∈ { (,•,) } from Eq (3) we have

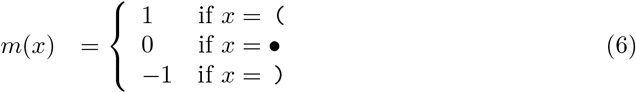

Define the distance *d*_0_(*x, y*) between characters *x, y* in the dot-bracket representation of a secondary structure by

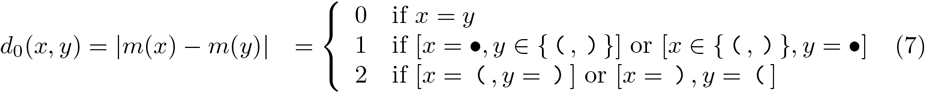

Let 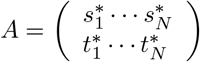 denote an alignment between two arbitrary secondary structures *s, t* of (possibly different) lengths *n, m*, where 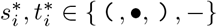 and – denotes the gap symbol. We define the *structural alignment distance* for *A* by summing 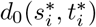 over those positions *i* where neither character 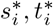 is a gap symbol, then adding *w*(*k*) for all size *k* gaps in *A*. Using previous definitions of incremental ensemble expected mountain height from Eq (3), we can generalize structural alignment distance from the simple case of comparing two dot-bracket representations of secondary structures to the more representative case of comparing the low-energy Boltzmann ensemble of secondary structures for RNA sequence **a** to that of RNA sequence **b**. Given sequences **a** = *a*_1_,…, *a_n_* and **b** = *b*_1_,…,*b_m_*, let 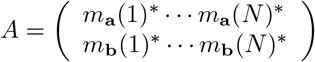 denote an alignment between the incremental ensemble expected mountain height *m*_a_(1) ··· *m*_a_(*n*) of **a** and and the ensemble incremental expected mountain height *m*_b_(1) ··· *m*_b_(*m*) of **b**. Generalize structural distance *d*_0_ defined in Eq (7) to *d*_1_ defined by *d*_1_(*a_i_*, *b_j_*) = |*m_a_*(*i*) – *m_b_*(*j*)|, where *m_a_*(*i*) and *m_b_*(*j*) are real numbers in the interval [–1, 1], and define *ensemble structural alignment distance* for *A* by summing *d*_1_(*a_i_*,*b_j_*) over all positions *i, j* for which neither character is a gap symbol, then adding positive weight *w*(*k*) for all size *k* gaps. By Eq (4) and Eq (5), it follows that an equivalent *ensemble structural similarity* measure between two positions *a_i_*, *b_j_*, denoted *STRSIM*(*a_i_, b_j_*), is obtained by multiplying *d*_1_ and *w*(*k*) by –1:

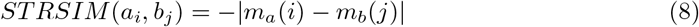

This equation will be used later, since our algorithm `RNAmountAlign` combines both sequence and ensemble structural similarity. Indeed, –|*m_a_*(*i*) – *m_b_*(*j*)| ∈ [–2, 0] with maximum value of 0 while RIB0SUM85-60, shown in Table 3, has similarity values in the interval [–1.86, 2.22]. In order to combine sequence with structural similarity, both ranges should be rendered comparable as shown in the next section.

### Pairwise alignment

In order to combine sequence and ensemble structural similarity, we determine a multiplicative scaling factor *α*_seq_ and an additive shift factor *α*_str_ such that the mean and standard deviation for the distribution of sequence similarity values from a RIBOSUM matrix [36] (after being multiplied by *α*_seq_) are equal to the mean and standard deviation for the distribution of structural similarity values from STRSIM (after additive shift of *α*_str_). The RIB0SUM85-60 nucleotide similarity matrix used in this paper is given in Table 3, and the distributions for RIBOSUM and STRSIM values are shown in Figure 2 for the 72 nt transfer RNA AL671879.2. Given query [resp. target] nucleotide frequencies 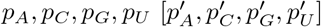 that sum to 1, the mean *μ*_seq_ and standard deviation *σ*_seq_ of RIBOSUM nucleotide similarities can be computed by

**Table 3.** RIB0SUM85-60 similarity matrix for RNA nucleotides from [36].

|  | A | C | G | U |
| --- | --- | --- | --- | --- |
| A | +2.22 | -1.86 | -1.46 | -1.39 |
| C | -1.86 | +1.16 | -2.48 | -1.05 |
| G | -1.46 | -2.48 | +1.03 | -1.74 |
| U | -1.39 | -1.05 | -1.74 | +1.65 |

**FIG. 2.**
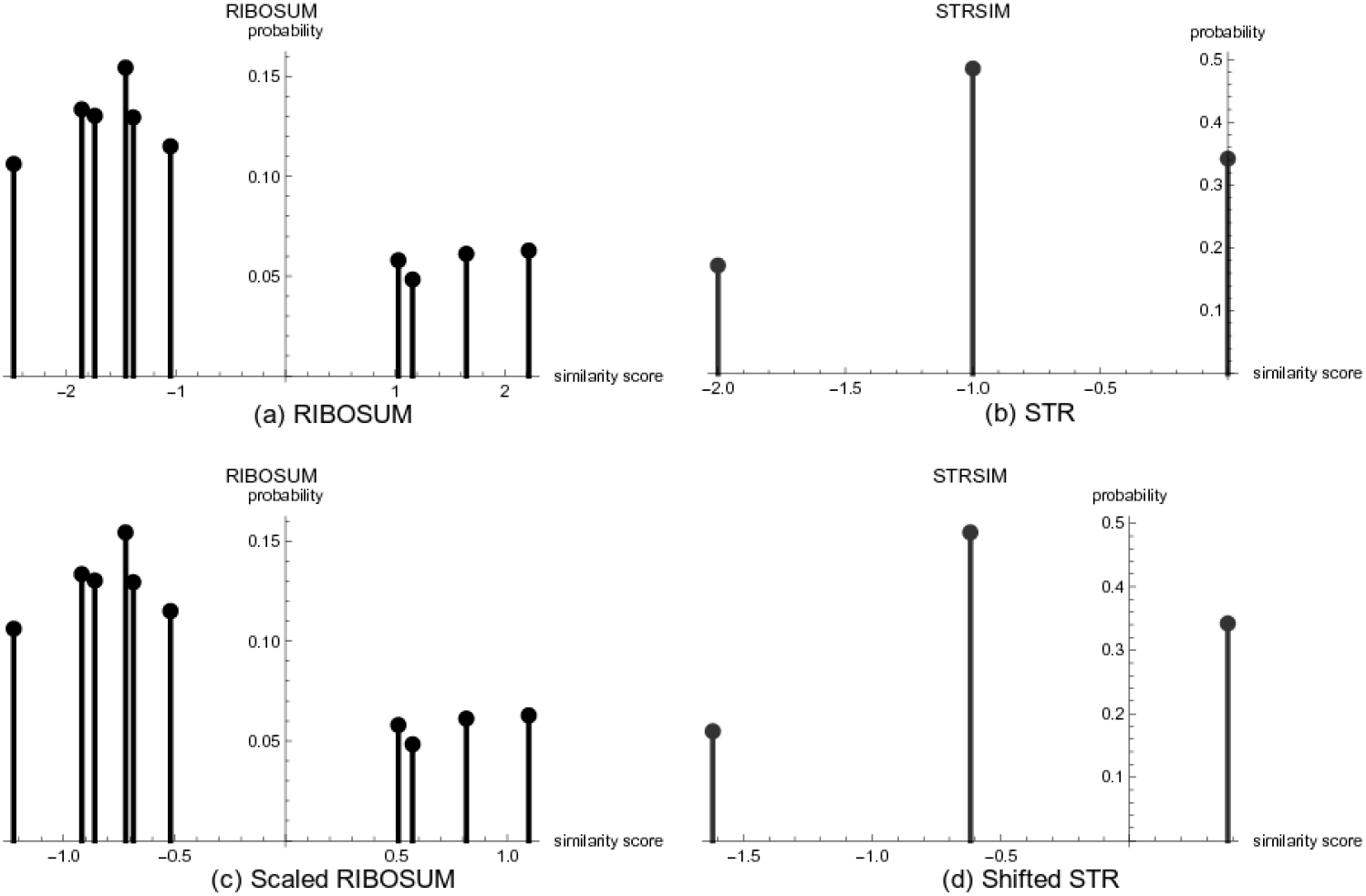
For 72 nt tRNA query sequence AL671879.2, nucleotide frequencies are approximately *p_A_* = 0.167, *p_C_* = 0.278, *p_G_* = 0.333, *p_U_* = 0.222, and for 69 nt tRNA target sequence D16498.1, nucleotide frequencies are approximately *p_A_* = 0.377, *p_C_* = 0.174, *p_G_* = 0.174, *p_U_* = 0.275. From the base pairing probabilities computed by `RNAfold -p`, we have query frequencies *p* ( = 0.3035, *p*• = 0.3930, *p*) = 0.3035 and target frequencies *p* ( = 0.2835, *p*• = 0.433, *p*) = 0.2835, so by Eqs (9,10,11,12), we have *μ*_seq_ = –0.9098, *σ*_seq_ = 1.4117 and *μ*_str_ = –0.8301, *σ*_str_ = 0.6968. By Eqs (13) and (14), we determine that RIBOSUM scaling factor *α*_seq_ = 0.4936 and *α*_str_ = 0.3810 (values shown only to 4-decimal places). Panels (a) resp. (b) show the distribution of RIBOSUM resp. STRSIM values for the nucleotide and base pairing probabilities determined from query and target, while panels (c) resp. (d) show the distribution of *α*_seq_-scaled RIBOSUM values resp. *α*_str_-shifted STRSIM values. It follows that distributions in panels (c) and (d) have the same (negative) mean and standard deviation.

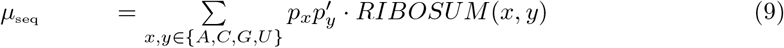

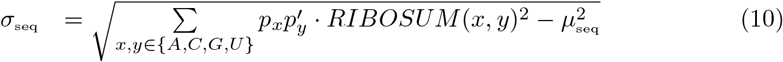

Setting *s*_0_(*x,y*) = –*d*_0_(*x,y*), where *d*_0_(*x,y*) is defined in Eq (7), for given query [resp. target] base pairing probabilities *p* (,*p*•,*p*) [resp. 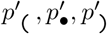] of dot-bracket characters, it follows that the mean *μ*_str_ and standard deviation *σ*_str_ of structural similarities can be computed by

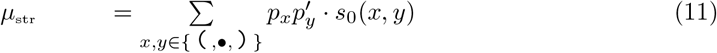

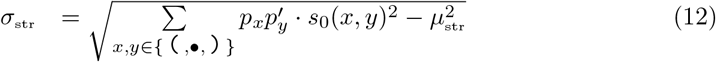

Now we compute a multiplicative factor *α*_seq_ and an additive shift term *α*_str_, both dependent on frequencies *p_A_,p_C_,p_G_,p_U_* and *p* (,*p*•,*p*), such that the mean [resp. standard deviation] of nucleotide similarity multiplied by *α*_seq_ is equal to the mean [resp. standard deviation] of structural similarity after addition of shift term *α*_str_:

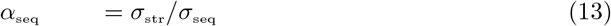

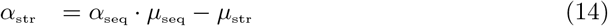

Given the query RNA **a** = *a*_1_,…,*a_n_* and target RNA **b** = *b*_1_,…,*b_m_* with incremental ensemble expected mountain heights *m*_a_(1) ··· *m*_a_(*m*) of **a**, *m*_b_(1) ··· *m*_b_(*m*) of **b**, and user-defined weight 0 ≤ *γ* ≤ 1, our final similarity measure is defined by

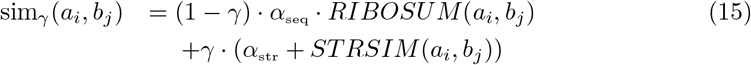

where *α*_seq_, *α*_str_ are computed by Eqs (13,14) depending on probabilities *p_A_,p_C_,p_G_,p_U_* [resp. 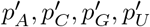] and *p* (,*p*•,*p*) [resp. 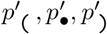] of the query [resp. target]. All benchmarking computations were carried out using *γ* = 1/2, although it is possible to use position-specific weight *γ_i,j_* defined as the average probability that *i* is paired in **a** and *j* is paired in **b**.

Our structural similarity measure is closely related to that of `STRAL`, which we discovered only after completing a preliminary version of this paper. Let 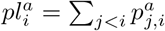 and 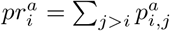 be the probability that position *i* of sequence a is paired to a position on the left or right, respectively. The similarity measure used in `STRAL` is defined by

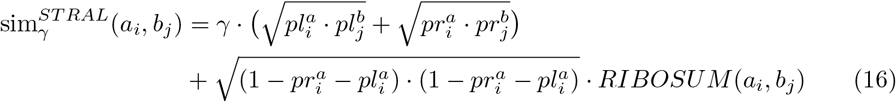

From Eq (15) and Eq (3) our measure can be defined as

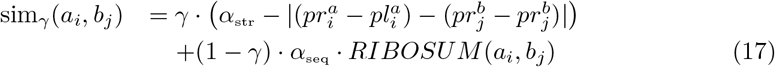

Though `RNAmountAlign` was developed independently much later than `STRAL`, our software offers functionalities unavailable in `STRAL`, which latter appears to be no longer maintained.^2^ For instance, `RNAmountAlign` supports local and semiglobal alignment, and reports *p*-values and E-values; these features are not available in `STRAL`.

To illustrate the method, suppose that the query [resp. target] sequence is the 72 nt tRNA AL671879.2 [resp. 69 nt tRNA D16498.1]. Then nucleotide query [resp. target] probabilities are (approximately) *p_A_* = 0.167, *p_C_* = 0.278, *p_G_* = 0.333, *p_U_* = 0.222, [resp. 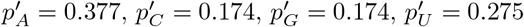]. From the base pairing probabilities returned by `RNAfold-p` [4], we determine that *p* ( = 0.3035, *p*• = 0.3930, *p*) = 0.3035 [resp. 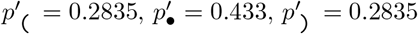]. Using these probabilities in Eqs (9-12), we determine that *μ*_seq_ = –0.9098, *σ*_seq_ = 1.4117, and *μ*_str_ = –0.8301, *σ*_str_ = 0.6968. By Eq (13) and Eq (14), we determine that RIBOSUM scaling factor *α*_seq_ = 0.4936 and *α*_str_ = 0.3810. It follows that the mean and standard deviation of *α*_seq_-scaled RIBOSUM values are identical with that of *α*_str_-shifted STRSIM values, hence can be combined in Eq (15). Since sequence identity of the `BRAliBase 2.1` alignment of these tRNAs is only 28%, we set structural similarity weight *γ* = 1 in Eq (15), and obtained a (perfect) global alignment computed by `RNAmountAlign`. Figure 2 depicts the distribution of RIBOSUM85-60 [resp. STRSIM] values in this case, both before and after application of scaling factor *α*_seq_ [resp. shift *α*_str_] – recall that *α*_seq_ and *α*_str_] depend on *p_A_,p_C_,p_G_,p_U_,p* (,*p*•,*p*) of tRNA AL671879.2 and 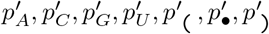 of tRNA D16498.1.

### Statistics for pairwise alignment

#### Karlin-Altschul statistics for local pairwise alignment

For a finite alphabet *A* and similarity measure *s*, suppose that the expected similarity 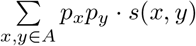 is negative and that *s*(*x, y*) is positive for at least one choice of *x, y*. In the case of `BLAST`, amino acid and nucleotide similarity scores are integers, for which the Karlin-Altschul algorithm was developed [12]. In contrast, `RNAmountAlign` similarity scores scores are not integers (or more generally values in a lattice), because Eq (15) combines real-valued *α*_seq_-scaled RIBOSUM nucleotide similarities with real-valued *α*_str_-shifted STRSIM structural similarities, which depend on query [resp. target] probabilities *p_A_,p_C_,p_G_,p_U_,p* (,*p*•,*p*) [resp. 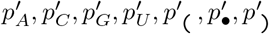]. For that reason, we use the following reformulation of a result by Karlin, Dembo and Kawabata [13], the similarity score *s*(*x, y*) for RNA nucleotides *x, y* is defined by Eq (15).

##### Theorem 1 (Theorem 1 of [13])

*Given similarity measure s between nucleotides in alphabet A* = {*A, C, G, U*}, *let λ*\* *be the unique positive root of* 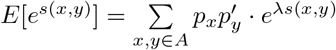, *and let random variable S_k_ denote the score of a length k gapless alignment. For large z*,

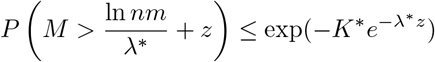

*where M denotes high maximal segment scores for local alignment of random RNA sequences a*_1_,…, *a_n_ and b*_1_,…, *b_m_, and where*

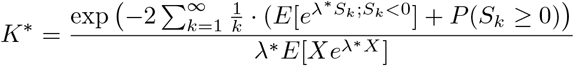

#### Fitting data to probability distributions

Data were fit to the normal distribution (ND) by the method of moments (i.e. mean and standard deviation were taken from data analysis). Data were fit to the extreme value distribution (EVD)

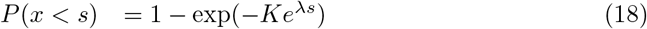

by an in-house implementation of maximum likelihood to determine *λ, K*, as described in supplementary information to [36]. Data were fit to the gamma distribution by using the function `fitdistr(x,’gamma’)` from the package `MASS` in the `R` programming language, which determines rate and shape parameters for the density function

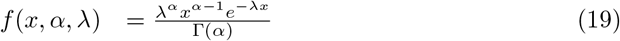

with where *α* is the shape parameter, the rate is 1/*λ*, where *λ* is known as the scale parameter.

### Multiple alignment

Suppose *p_A_,p_C_,p_G_,p_U_* are the nucleotide probabilities obtained after the concatenation of all sequences. Let *p* (,*p*•,*p*) be computed by individually folding each sequence and taking the arithmetic average of probabilities of (, • and) over all sequences. The mean and standard deviation of sequence and structure similarity are computed similar to Eqs (9-12).

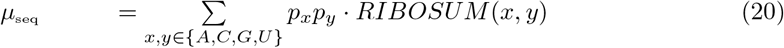

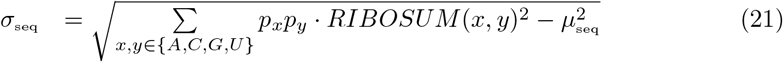

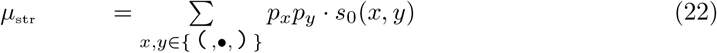

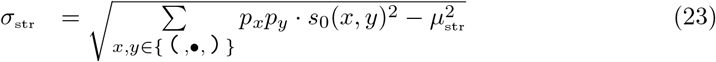

Sequence multiplicative scaling factor *α*_seq_ and the structure additive shift factor *α*_str_ are computed from these values using Eqs (13,14).

`RNAmountAlign` implements progressive multiple alignment using `UPGMA` to construct the guide tree. In `UPGMA`, one first defines a similarity matrix *S*, where *S*[*i, j*] is equal to (maximum) pairwise sequence similarity of sequences *i* and *j*. A rooted tree is then constructed by progressively creating a parent node of the two closest siblings. Parent nodes are profiles (PSSMs) that represent alignments of two or more sequences, hence can be treated as pseudo-sequences in a straightforward adaptation of pairwise alignment to the alignment of profiles. Let’s consider an alignment of *N* sequences 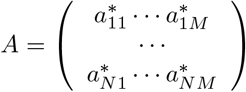 composed of *M* columns. Let 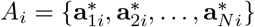 denote column *i* of the alignment (for 1 ≤ *i* ≤ *M*). Suppose *p*(*i, x*), for *x* ∈ {*A, C, G, U*, –}, indicates the probability of occurrence of a nucleotide or gap at column *i* of alignment *A*. Then sequence similarity SEQSIM between two columns is defined by

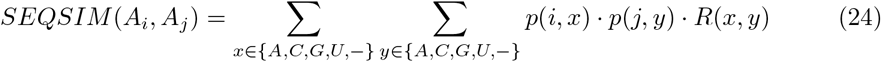

where

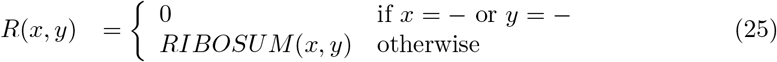

The structural measure for a profile is computed from the incremental ensemble heights averaged over each column. Let *mA*(*i*) denote the arithmetic average of incremental ensemble mountain height at column *A_i_*

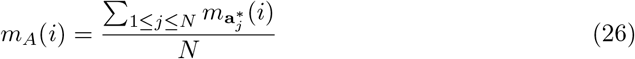

where 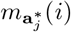 is the incremental ensemble mountain height at position *i* of sequence 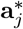 obtained from Eq (3). Here, let 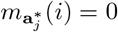 if 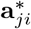 is a gap. Structural similarity between two columns is defined by

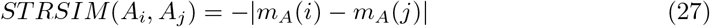

Finally, the combined sequence/structure similarity is computed from

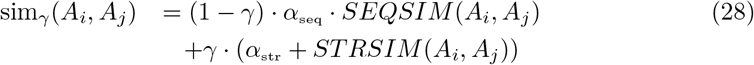

### Benchmarking

#### Accuracy measures

Sensitivity, positive predictive value, and F1-measure for pairwise alignments were computed as follows. Let 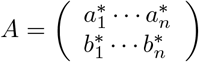 denotes an alignment, where *a_i_, b_i_* ∈ {*A, C, G, U*, –}, and the aligned sequences include may contain gap symbols – provided that it is not the case that both 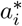 and 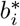 are gaps. The number TP of true positives [resp. FP of false positives] is the number of alignment pairs 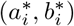 in the predicted alignment that belong to [resp. do not belong to] the reference alignment. The sensitivity (*Sen*) [resp. positive predictive value (*PPV*)] of a predicted alignment is TP divided by reference alignment length [resp. TP divided by predicted alignment length]. The *F*1-score is the harmonic mean of sensitivity and PPV, so 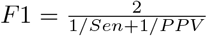. For the computation of *Sen, PPV*, and *F*1, pairs of the form (*X*, —) and (—, *X*) are also counted. In the case of local alignment, since the size of the reference alignment is unknown, only the predicted alignment length and PPV are reported. To compute the accuracy of multiple alignment, we used sum-of-pair-scores (SPS) [11], defined as follows. Suppose that *A* denotes a multiple alignment of the form 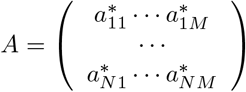. For 1 ≤ *i,j* ≤ *M*, 1 ≤ *k* ≤ *N* define *p_ijk_* = 1 if 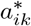 is aligned with 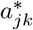 in both the reference and predicted alignments, and *p_ijk_* = 0 otherwise. Sum-of-pairs score SPS is then the sum, taken over all *i, j, k*, of the *p_ijk_*. Though SPS can be considered as the average sensitivity, taken over all sequence pairs in the alignment, this is not technically the case, since our definition of sensitivity also counts pairs of the form (*X*, —) and (—, *X*) from the reference alignment.

To measure the conservation of secondary structures in alignments, structural conservation index (SCI) was computed using `RNAalifold` [37]. *RNAalifold* computes SCI as the ratio of the free energy of the alignment, computed by `RNAalifold`, with the average minimum free energy of individual structures in the alignment. SCI values close to 1 [resp. 0] indicate high [resp. low] structural conservation. All computations made with Vienna RNA Package used version 2.1.7 [4] using default Turner 2004 energy parameters [38]).

#### Dataset for global and local alignment comparison

For *pairwise global* alignment benchmarking in Table 4 and Figures 3, 4, S2 and S3 all 8976 pairwise alignments in k2 from `BRAliBase 2.1` database [33] were used. For *multiple global* alignment benchmarking in Fig 7, k5 `BRAliBase 3` was used [39]. This dataset includes 583 reference alignments, each composed of 5 sequences. For *pairwise local alignment* benchmarking, 75 pairwise alignments having sequence identity ≤ 70% were randomly selected from each of 20 well-known families from the Rfam 12.0 database [40], many of which were considered in a previous study [41], yielding a total of 1500 alignments. Following [42], these alignments were trimmed on the left and right, so that both first and last aligned pairs of the alignment do not contain a gap symbol. For sequences **a** = *a*_1_,…, *a_n_* [resp. **b** = *b*_1_,…, *b_m_*] from each alignment, random sequences **a**’ [resp. **b**’] were generated with the same nucleotide frequencies, then a random position was chosen in **a**’ [resp. **b**’] in which to insert **a** [resp. **b**], thus resulting in a pair of sequences of lengths 4*n* and 4*m*. Finally, since sequence identity was at most 70%, the RIBOSUM70-25 similarity matrix was used in `RNAmountAlign`. Preparation of the benchmarking dataset for local alignment was analogous to the method used in *multiple* local alignment of [42]. We used `LocARNA` (version 1.8.7), `FOLDALIGN` (version 2.5), `LARA` (version 1.3.2) `DYNALIGN` (from version 5.7 of `RNAstructure`), and `STRAL` (in-house implementation due to unavailability) for benchmarking.

**Table 4.** Average F1 scores (± one standard deviation) for *pairwise global alignment* of `RNAmountAlign` and four widely used RNA sequence/structure alignment algorithms on the benchmarking set of 8976 pairwise alignments from the `BRaliBase K2` database [33]. For each indicated Rfam family, the the number of alignments (NumAln), sequence identity (SeqId), and F1-scores for `RNAmountAlign, LocARNA, LARA, FOLDALIGN`, and `DYNALIGN` are listed, along with pooled averages over all 8976 pairwise alignments. Parameters used in Eq (15) for `RNAmountAlign` were similarity matrix RIBOSUM85-60, structural similarity weight *γ* = 1/2, gap initiation *g_i_* = –3, gap extension *g_e_* = –1.

| Type | NumAln | SeqId | MA(F) | LocARNA(F) | LARA(F) | FA(F) | DA(F) | STRAL(F) |
| --- | --- | --- | --- | --- | --- | --- | --- | --- |
| 5.8S rRNA | 76 | 0.72 ± 0.13 | 0.90 ± 0.09 | 0.82 ± 0.07 | 0.87 ± 0.15 | 0.89 ± 0.11 | 0.66 ± 0.22 | 0.88 ± 0.12 |
| 5S rRNA | 1162 | 0.60 ± 0.14 | 0.84 ± 0.16 | 0.87 ± 0.13 | 0.85 ± 0.16 | 0.86 ± 0.14 | 0.69 ± 0.17 | 0.82 ± 0.20 |
| Cobalamin | 188 | 0.43 ± 0.10 | 0.56 ± 0.16 | 0.38 ± 0.17 | 0.49 ± 0.20 | 0.43 ± 0.24 | 0.36 ± 0.19 | 0.54 ± 0.17 |
| Entero 5 CRE | 48 | 0.88 ± 0.06 | 0.98 ± 0.04 | 0.99 ± 0.04 | 0.99 ± 0.05 | 0.99 ± 0.02 | 0.87 ± 0.13 | 0.97 ± 0.06 |
| Entero CRE | 65 | 0.80 ± 0.07 | 1.00 ± 0.00 | 0.99 ± 0.03 | 0.96 ± 0.07 | 0.99 ± 0.04 | 0.76 ± 0.17 | 1.00 ± 0.03 |
| Entero OriR | 49 | 0.84 ± 0.06 | 0.95 ± 0.07 | 0.92 ± 0.09 | 0.94 ± 0.08 | 0.94 ± 0.07 | 0.84 ± 0.15 | 0.95 ± 0.07 |
| gcvT | 167 | 0.44 ± 0.13 | 0.61 ± 0.19 | 0.61 ± 0.24 | 0.57 ± 0.25 | 0.40 ± 0.33 | 0.44 ± 0.19 | 0.62 ± 0.20 |
| Hammerhead 1 | 53 | 0.71 ± 0.17 | 0.89 ± 0.13 | 0.90 ± 0.11 | 0.87 ± 0.16 | 0.83 ± 0.25 | 0.52 ± 0.27 | 0.88 ± 0.16 |
| Hammerhead 3 | 126 | 0.66 ± 0.21 | 0.86 ± 0.20 | 0.88 ± 0.21 | 0.88 ± 0.20 | 0.80 ± 0.31 | 0.71 ± 0.31 | 0.90 ± 0.16 |
| HCV SLIV | 98 | 0.85 ± 0.05 | 0.99 ± 0.03 | 0.98 ± 0.04 | 0.98 ± 0.03 | 0.99 ± 0.03 | 0.81 ± 0.34 | 0.99 ± 0.03 |
| HCV SLVII | 51 | 0.83 ± 0.09 | 0.97 ± 0.06 | 0.96 ± 0.06 | 0.93 ± 0.10 | 0.95 ± 0.07 | 0.71 ± 0.22 | 0.95 ± 0.07 |
| HepC CRE | 45 | 0.86 ± 0.06 | 1.00 ± 0.00 | 1.00 ± 0.00 | 1.00 ± 0.00 | 1.00 ± 0.00 | 0.77 ± 0.29 | 1.00 ± 0.00 |
| Histone3 | 84 | 0.78 ± 0.09 | 1.00 ± 0.00 | 1.00 ± 0.00 | 1.00 ± 0.00 | 1.00 ± 0.00 | 1.00 ± 0.00 | 1.00 ± 0.00 |
| HIV FE | 733 | 0.87 ± 0.04 | 1.00 ± 0.02 | 1.00 ± 0.02 | 0.98 ± 0.05 | 0.99 ± 0.05 | 0.64 ± 0.29 | 1.00 ± 0.02 |
| HIV GSL3 | 786 | 0.86 ± 0.04 | 0.99 ± 0.02 | 0.99 ± 0.02 | 0.98 ± 0.05 | 0.99 ± 0.02 | 0.80 ± 0.19 | 0.99 ± 0.02 |
| HIV PBS | 188 | 0.92 ± 0.02 | 1.00 ± 0.01 | 1.00 ± 0.01 | 1.00 ± 0.02 | 0.99 ± 0.03 | 0.91 ± 0.11 | 1.00 ± 0.01 |
| Intron gpII | 181 | 0.46 ± 0.13 | 0.64 ± 0.17 | 0.64 ± 0.17 | 0.63 ± 0.17 | 0.50 ± 0.28 | 0.49 ± 0.18 | 0.65 ± 0.15 |
| IRES HCV | 764 | 0.65 ± 0.11 | 0.88 ± 0.16 | 0.45 ± 0.19 | 0.86 ± 0.17 | 0.68 ± 0.38 | 0.85 ± 0.08 | 0.88 ± 0.08 |
| IRES Picorna | 181 | 0.84 ± 0.07 | 0.97 ± 0.03 | 0.61 ± 0.04 | 0.96 ± 0.04 | 0.95 ± 0.04 | 0.85 ± 0.11 | 0.96 ± 0.04 |
| K chan RES | 124 | 0.74 ± 0.10 | 0.99 ± 0.02 | 0.98 ± 0.05 | 0.89 ± 0.19 | 0.95 ± 0.08 | 0.58 ± 0.26 | 0.95 ± 0.11 |
| Lysine | 80 | 0.50 ± 0.13 | 0.72 ± 0.13 | 0.54 ± 0.15 | 0.71 ± 0.18 | 0.66 ± 0.16 | 0.50 ± 0.16 | 0.72 ± 0.15 |
| Retroviral psi | 89 | 0.88 ± 0.03 | 0.93 ± 0.03 | 0.93 ± 0.03 | 0.93 ± 0.03 | 0.92 ± 0.04 | 0.74 ± 0.12 | 0.93 ± 0.04 |
| S box | 91 | 0.60 ± 0.10 | 0.75 ± 0.13 | 0.76 ± 0.16 | 0.79 ± 0.14 | 0.67 ± 0.24 | 0.54 ± 0.16 | 0.77 ± 0.12 |
| SECIS | 114 | 0.44 ± 0.16 | 0.59 ± 0.21 | 0.62 ± 0.21 | 0.57 ± 0.25 | 0.54 ± 0.25 | 0.39 ± 0.24 | 0.61 ± 0.20 |
| sno 14q I II | 44 | 0.75 ± 0.10 | 0.92 ± 0.10 | 0.89 ± 0.16 | 0.85 ± 0.20 | 0.89 ± 0.19 | 0.58 ± 0.27 | 0.91 ± 0.13 |
| SRP bact | 114 | 0.48 ± 0.16 | 0.65 ± 0.21 | 0.66 ± 0.21 | 0.63 ± 0.25 | 0.65 ± 0.21 | 0.51 ± 0.22 | 0.61 ± 0.25 |
| SRP euk arch | 122 | 0.51 ± 0.20 | 0.62 ± 0.29 | 0.35 ± 0.17 | 0.64 ± 0.28 | 0.64 ± 0.26 | 0.50 ± 0.26 | 0.61 ± 0.29 |
| T-box | 18 | 0.68 ± 0.15 | 0.77 ± 0.17 | 0.49 ± 0.17 | 0.68 ± 0.25 | 0.70 ± 0.17 | 0.59 ± 0.21 | 0.74 ± 0.15 |
| TAR | 286 | 0.87 ± 0.04 | 0.99 ± 0.03 | 0.99 ± 0.02 | 0.99 ± 0.03 | 0.98 ± 0.04 | 0.83 ± 0.19 | 0.99 ± 0.04 |
| THI | 321 | 0.45 ± 0.10 | 0.68 ± 0.16 | 0.66 ± 0.20 | 0.68 ± 0.18 | 0.50 ± 0.29 | 0.48 ± 0.18 | 0.65 ± 0.20 |
| tRNA | 2039 | 0.43 ± 0.12 | 0.75 ± 0.21 | 0.85 ± 0.16 | 0.82 ± 0.19 | 0.76 ± 0.27 | 0.66 ± 0.23 | 0.72 ± 0.22 |
| U1 | 82 | 0.63 ± 0.17 | 0.79 ± 0.17 | 0.70 ± 0.13 | 0.79 ± 0.19 | 0.80 ± 0.14 | 0.67 ± 0.20 | 0.77 ± 0.17 |
| U2 | 112 | 0.64 ± 0.16 | 0.75 ± 0.17 | 0.63 ± 0.13 | 0.76 ± 0.19 | 0.73 ± 0.22 | 0.59 ± 0.19 | 0.75 ± 0.18 |
| U6 | 30 | 0.83 ± 0.06 | 0.93 ± 0.05 | 0.89 ± 0.09 | 0.90 ± 0.08 | 0.88 ± 0.10 | 0.72 ± 0.14 | 0.93 ± 0.06 |
| UnaL2 | 138 | 0.77 ± 0.08 | 0.93 ± 0.08 | 0.92 ± 0.09 | 0.89 ± 0.15 | 0.91 ± 0.10 | 0.65 ± 0.29 | 0.94 ± 0.08 |
| yybP-ykoY | 127 | 0.39 ± 0.14 | 0.58 ± 0.20 | 0.54 ± 0.23 | 0.57 ± 0.25 | 0.40 ± 0.33 | 0.46 ± 0.22 | 0.56 ± 0.20 |
| Pooled Average | 249.33 | 0.63 | 0.84 | 0.81 | 0.84 | 0.8 | 0.68 | 0.82 |

**Fig 3.**
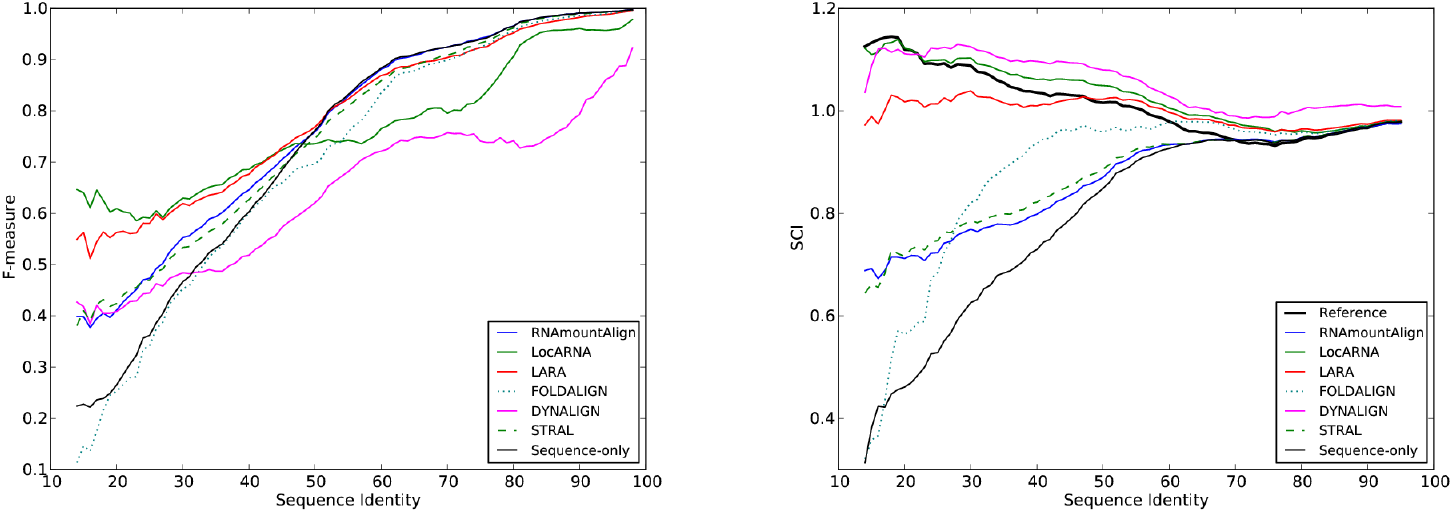
F1-measure (Left) and structural conservation index (SCI) (Right) for *pairwise global alignments* using `RNAmountAlign, LocARNA, LARA, FOLDALIGN, DYNALIGN, STRAL` and sequence-only(*γ* = 0). F1-measure and SCI are shown as a function of alignment sequence identity for pairwise alignments in the `BRAliBase 2.1` database used for benchmarking.

#### Dataset for correlation of *p*-values for different distribution fits

A pool of 2220 sequences from the Rfam 12.0 database [40] was created as follows. One sequence was selected from each Rfam family having average sequence length at most 200 nt, with the property that the base pair distance between its minimum free energy (MFE) structure and the Rfam consensus structure was a minimum. Subsequently, for each of 500 randomly selected *query* sequences from the pool of 2220 sequences, 1000 random *target* sequences of length 400 nt were generated to have the same expected nucleotide frequency as that of the query. For each query and random target, five semiglobal (query search) alignments were created using gap initiation costs of *g_i_* ∈ {–1, –2, –3, –4, –5} with gap extension cost *g_e_* equal to one-third the gap initiation cost. For each alignment score *x* for query and random target, the *p*-value was computed as 1 – *CDF*(*x*) for ND, EVD and GD, where *CDF*(*x*) is the cumulative density function evaluated at *x*. Additionally, a heuristic *p*-value was determined by calculating the proportion of alignment scores for given query that exceed *x*.

## Results

We benchmarked `RNAmountAlign`’s performance for pairwise and multiple alignments on `BraliBase k2` and k5 datasets, respectively.

### Pairwise alignment

Figures 3, S2 and S3 depict running averages of *pairwise global alignment* F1-measure, sensitivity, and positive predictive value (PPV) for the software described in this paper, as well as for `LocARNA, FOLDALIGN, LARA, DYNALIGN`, and `STRAL`. For pairwise benchmarking, reference alignments of size 2, a.k.a. K2, were taken from the `BRAliBase 2.1` database [33]. `BRAliBase 2.1 K2` data are based on seed alignments of the Rfam 7.0 database, and consist of 8976 alignments of RNA sequences from 36 Rfam families.

Running averages of sensitivity, positive predictive value, and F1-measure, averaging over windows of size 11 nt (interval [*k* – 5, *k* + 5]), were computed as a function of sequence identity, where it should be noted that the number of pairwise alignments for different values of sequence identity can vary for the `BRAliBase 2.1` data (e.g. there are only 35 pairwise alignments having sequence identity < 20%). Default parameters were used for all other software. For our software `RNAmountAlign`, gap initiation cost was −3, gap extension −1, and sequence/structure weighting parameter *γ* was 0.5 (value obtained by optimizing on a small set of 300 random alignments from Rfam 12.0, not considered in training or testing set). The sequence-only alignment is computed from `RNAmountAlign` with the same gap penalties, but for *γ* = 0. While its accuracy is high, `RNAmountAlign` is faster by an order of magnitude than `LocARNA, LARA, FOLDALIGN`, and `DYNALIGN` – indeed, algorithmic time complexity of our method is *O*(*n*^3^) compared with *O*(*n*^4^) for these methods. Since `STRAL` could not be compiled on any of our systems, we implemented its algorithm by modifying `RNAmountAlign` and obtained results for `STRAL`’s default parameter settings. Therefore, the run time of `STRAL` is identical to `RNAmountAlign` but we achieve slightly higher F1-measure, sensitivity and PPV. Moreover, `RNAmountAlign` supports semiglobal and local alignments as well as reporting *p*-values. The right panel of Fig 4 depicts actual run times of the fastest software, `RNAmountAlign`, with the next fastest software, `LARA`. Unlike the graph in the left panel, actual run times are shown, graphed as a function of sequence length, rather than logarithms of moving averages.

**Fig 4.**
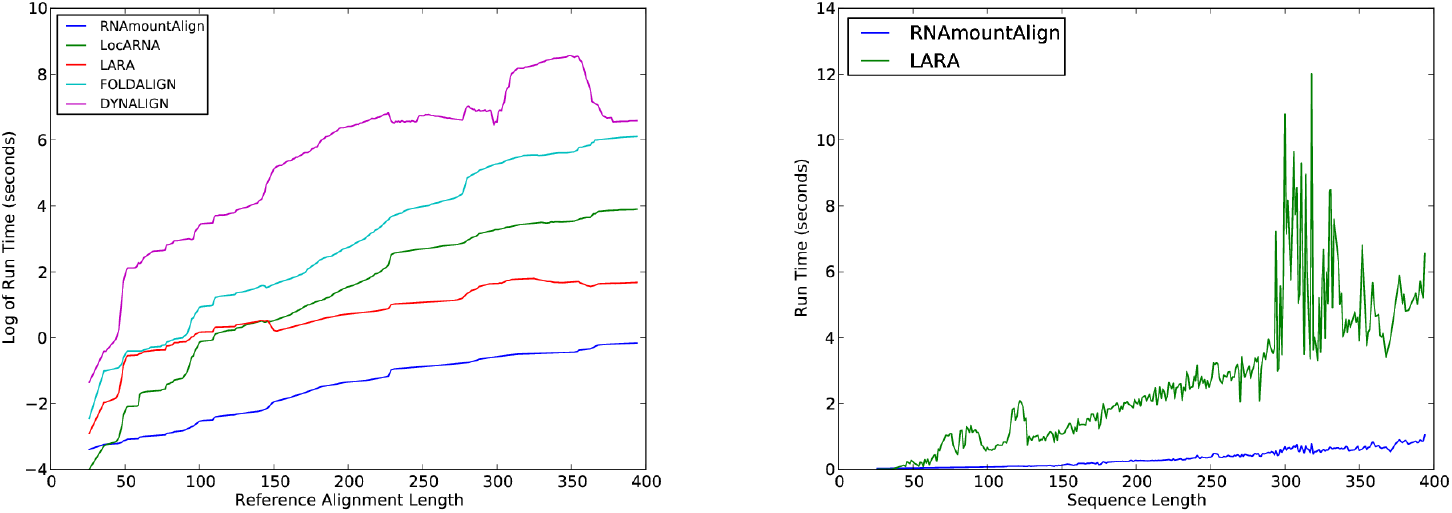
Run time of *pairwise global alignment* for `RNAmountAlign, LocARNA, LARA, FOLDALIGN`, and `DYNALIGN`. (Left) Log run time is shown as a function of seed length for pairwise alignments in the `BRAliBase 2.1` database used for benchmarking. Window size of 51 is used for the computation of moving average. (Right) Actual run time for `RNAmountAlign` and `LARA` on the same data. Unlike the left panel the actual run time is shown, rather than log run time, without any moving average taken.

In addition, Table 4 displays average pairwise global alignment F1 scores for `RNAmountAlign, LocARNA, LARA, FOLDALIGN, DYNALIGN`, and `STRAL` when benchmarked on 36 families from the `BRaliBase K2` database comprising altogether 8976 RNA sequences with average length of 249.33. Averaging over all sequences, the F1 scores for the programs just mentioned were respectively 0.8370, 0.7808, 0.8406, 0.7977, 0.6822, 0.8247; i.e. F1 score 0.8406 of `LARA` slightly exceeded the F1 score 0.8370 of `RNAmountAlign` and 0.8247 of `STRAL`, while other methods trailed by several percentage points. Supplementary Information (SI) Tables S1 and S2 display values for global alignment sensitivity and positive predictive value, benchmarked on the same data for the same programs – these results are similar to the F1-scores in Tables 2 and 4.

Although there appears to be no universally accepted criterion for quality of local alignments, Table 5 shows pairwise local alignment comparisons for the above-mentioned methods supporting local alignment: `RNAmountAlign, FOLDALIGN`, and `LocARNA`. We had intended to include `SCARNA_LM` [42] in the benchmarking of multiple local alignment software; however, `SCARNA_LM` no longer appears to be maintained, since the web server is no longer functional and no response came from our request for the source code. Since the reference alignments for the local benchmarking dataset are not known, and sensitivity depends upon the length of the reference alignment, we only report local alignment length and positive predictive value. Abbreviating `RNAmountAlign` by MA, `FOLDALIGN` by FA, and `LocARNA` by LOC, Table 5 shows average run time in seconds of MA (2.30 ± 2.12), FA (625.53 ± 2554.61), LOC (5317.96 ± 8585.19), average alignment length of reference alignments (118.67 ± 47.86), MA (50.35 ± 42.33), FA (114.86 ± 125.33), LOC (556.82 ± 227.00), and average PPV scores MA (0.53 ± 0.42), FA (0.64 ± 0.36), LOC (0.03 ± 0.04).

**Table 5.**
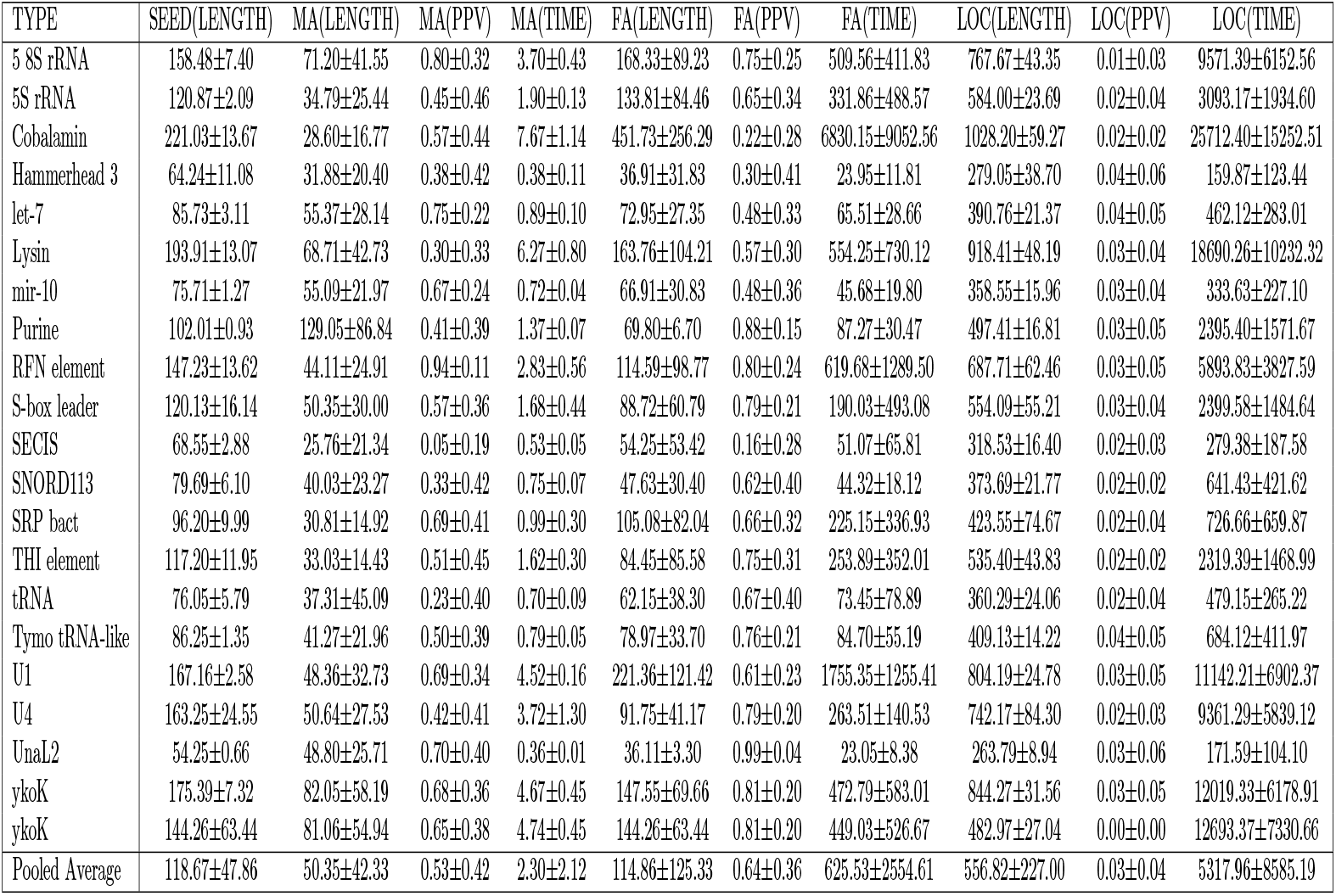
Comparison of alignment length and positive predictive value (PPV) for *pairwise local alignment* by `RNAmountAlign` against the widely used local alignment software `FOLDALIGN` and `LocARNA`. Local alignment benchmarking was performed on 1500 pairwise alignments (75 alignments per family, 20 Rfam families) extracted from the Rfam 12.0 database [40], and prepared in a manner analogous to that of the dataset used in benchmarking *multiple* local alignment in [42] – see text for details. Parameters used in Eq (15) of the main text for `RNAmountAlign` were structural similarity weight *γ* = 1/2, gap initiation *g_i_* = –3, gap extension *g_e_* = –1; since reference alignments were required to have at most 70% sequence identity, nucleotide similarity matrix RIBOSUM8570-25 was used in `RNAmountAlign`.

Taken together, these results suggest that `RNAmountAlign` has comparable accuracy, but much faster run time, hence making it a potentially useful tool for genome scanning applications. Here it should be stressed that all benchmarking results used equally weighted contributions of sequence and ensemble structural similarity; i.e. parameter *γ* =1/2 when computing similarity by Eq (15). By setting *γ* =1, `RNAmountAlign` alignments depend wholly on structural similarity (see Figure 1). Indeed, for the following `BRAliBase 2.1` alignment with 28% sequence identity, by setting *γ* =1, `RNAmountAlign` returns the correct alignment.

~~~
GGGGAUGUAGCUCAGUGGUAGAGCGCAUGCUUCGCAUGUAUGAGGCCCCGGGUUCGAUCCCCGGCAUCUCCA
GUUUCAUGAGUAUAGC---AGUACAUUCGGCUUCCAACCGAAAGGUUUUUGUAAACAACCAAAAAUGAAAUA
~~~

of 72 nt tRNA AL671879.2 with 69 nt tRNA D16387.1. Fig 1 shows the superimposed mountain heights for this alignment.

### Statistics for pairwise alignment

Fig 5 shows fits of the relative frequency histogram of alignment scores with the normal (ND), extreme value (EVD) and gamma (GD) distributions, where local [resp. semiglobal] alignment scores are shown in the left [resp. right] panel. The EVD provides the best fit for local alignment sequence-structure similarity scores, as expected by Karlin-Altschul theo [12,13]. Moreover, Fig 6 shows a 96% correlation between (expect) E-values computed by our implementation of the Karlin-Altschul method, and E-values obtained by maximum likelihood fitting of local alignment scores. In contrast, the ND provides the best fit for semiglobal sequence/structure alignment similarity scores, at least for the sequence considered in Fig 5. This is not an isolated phenomenon, as shown in Fig 6, which depicts scatter plots, Pearson correlation values and sums of squared residuals (SSRs) when computing *p*-values for semiglobal (query search) alignment scores between Rfam sequences and random RNA. As explained earlier, a pool of 2220 sequences from the Rfam 12.0 database [40] was created by selecting one sequence of length at most 200 nt from each family, with the property that base pair distance between its minimum free energy (MFE) structure and the Rfam consensus structure was a minimum. Then 500 sequences were randomly selected from this pool, and for each of five gap initiation and extension costs *g_i_* = –5, –4, –3, –2, –1 with 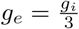. Taking each of the 500 sequences successively as query sequence and for each choice of parameters, 1000 random 400 nt RNAs were generated with the same expected nucleotide relative frequency as that of the query. For each alignment score *z* for query and random target, the *p*-value was computed as 1 minus the cumulative density function, 1 – *CDF*(*z*), for fitted normal (ND), extreme value (EVD) and gamma (GD) distributions, thus defining 1000 *p*-values. Additionally, a heuristic *p*-value was determined by calculating the proportion of alignment scores for given query that exceed *z*. For each set of 2.5 million (500 × 5 × 1000) *p*-values (heuristic, ND, EVD, GD), Pearson correlation values were computed and displayed in the upper triangular portion of Fig 6, with SSRs shown in parentheses. Note that residuals were computed for regression equation row = *m* · column + *b*, where column values constitute the independent variable. Assuming that heuristic *p*-values constitute the reference standard, it follows that *p*-values computed from the normal distribution correlate best with semiglobal alignment scores computed by `RNAmountAlign`.

**Fig 5.**
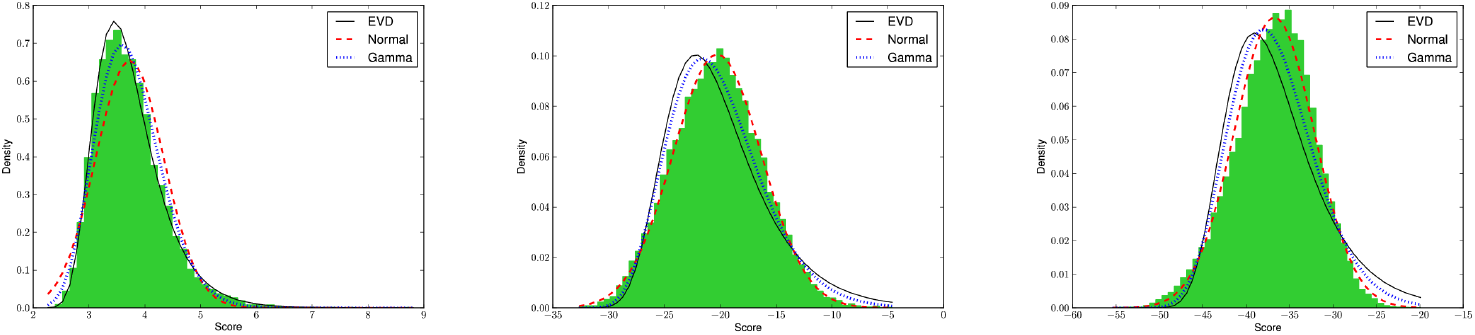
Fits of 30-bin relative frequency histograms of scores for *local* (left), *semiglobal* (middle) and *global* (right) alignments produced by `RNAmountAlign` for the randomly chosen 5S rRNA AY544430.1:375-465 from Rfam 12.0 database having A,C,G,U relative frequency of 0.25, 0.27, 0.26, 0.21. A total of 10,000 random sequences having identical expected nucleotide relative frequencies were generated, each of length 400 nt for local/semiglobal and 100 nt for global. Local (left), semiglobal (middle) and global (right) alignments were computed by `RNAmountAlign`, in each case fitting the data with the normal (ND), extreme value (EVD) and gamma (GD) distributions. As expected by Karlin-Altschul theory [12], local alignment scores are best fit by EVD, while semiglobal alignment scores are best fit by ND (results supported by data not shown, involving computations of variation distance, symmetrized Kullback-Leibler distance, and *χ*^2^ goodness-of-fit tests).

**Fig 6.**
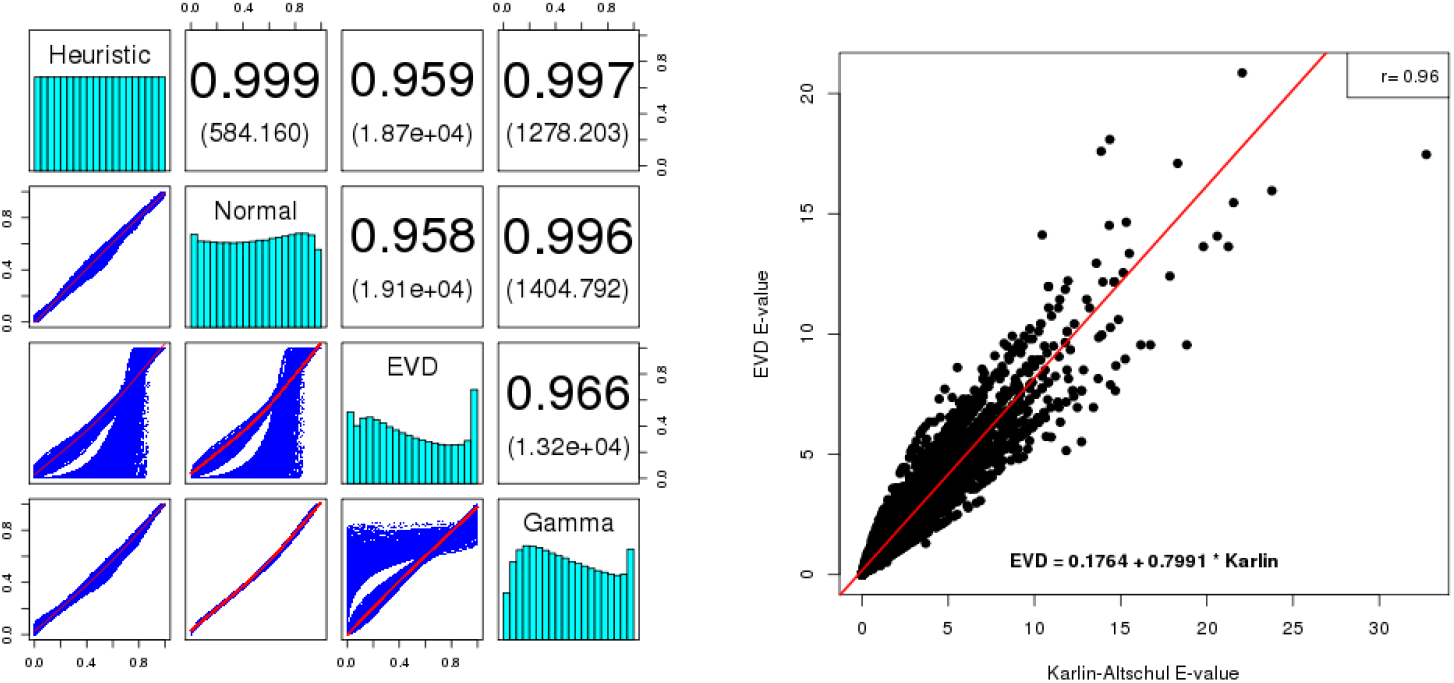
(**Left**)Pearson correlation values and scatter plots for *p*-values of *semiglobal alignment* (query search) scores between Rfam sequences and random RNA. For each score in a set of 2.5 million global pairwise alignment scores, a *p*-value was computed by direct counts (heuristic), or by data fitting the normal (ND), extreme value (EVD), or gamma (GD) distributions. Pairwise Pearson correlation values were computed and displayed in the upper triangular portion of the figure, with sums of squared residuals shown in parentheses, and histograms of *p*-values along the diagonal. It follows that ND *p*-values correlate best with heuristic *p*-values, where the latter is assumed to be the gold standard. (**Right**)Scatter plot of expect values *E*_ML_, computed by maximum likelihood, following the method described in [36] (*y*-axis) and expect values *E*_KA_, computed by our implementation of the Karlin-Altschul, as described in the text. The regression equation is *E*_ML_ = 0.1764 + 0.7991 · *E*_KA_; Pearson correlation between *E*_ML_ and *E*_KA_ is 96%, with correlation *p*-value of 2 · 10^−16^. Expect values were determined from local alignment scores computed by the genome scanning form of `RNAmountAlign` with query tRNA AB031215.1/9125-9195 and targets consisting of 300 nt windows (with 200 nt overlap) from *E. coli str. K-12* substr. MG1655 with GenBank accession code AKVX01000001.1. From the tRNA query sequence, the values *p_A_,p_C_,p_G_,p_U_* for nucleotide relative frequencies, are determined, then average base pairing probabilities *p* (,*p*•,*p*) are computed by RNAfold -p [4]. For the current 300 nt target window, the nucleotide relative frequencies 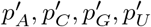 are computed, then precomputed probabilities 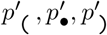 are obtained from SI Table S3. From these values, scaling factor *α*_seq_ and shift *α*_str_, were computed; with structural similarity weight *γ* = 1/2, the overall similarity function from Eq (15) in the text was determined.

Earlier studies have suggested that protein global alignment similarity scores using PAM120, PAM250, BLOSUM50, and BLOSUM62 matrices appear to be fit best by the gamma distribution (GD) [43], and that semiglobal RNA sequence alignment similarity scores (with no contribution from structure) appear to be best fit by GD [44]. However, in our preliminary studies (not shown), it appears that the type of distribution (ND, EVD, GD) that best fits `RNAmountAlign` semiglobal alignment depends on the gap costs applied (indeed, for certain choices, EVD provides the best fit). Since there is no mathematical theory concerning alignment score distribution for global or semiglobal alignments, it must be up to the user to decide which distribution provides the most reasonable *p*-values.

### Multiple alignment

We benchmarked `RNAmountAlign` with the software `LARA, mLocARNA, FOLDALIGNM` and `Multilign` for *multiple global* K5 alignments in `Bralibase 3. STRAL` is not included since the source code could not be compiled. Fig 7 indicates average SPS and SCI as a function of average pairwise sequence identity (APSI). We used the `-sci` flag of `RNAalifold` to compute SCI from the output of each software without reference to the reference alignment. Fig 7 indicates that SCI values for outputs from various alignment algorithms is higher than the SCI value from reference alignments, suggesting that the consensus structure obtained from sequence/structure alignment algorithms has a larger number of base pairs than the the consensus structure obtained from reference alignments (this phenomenon was also in [45]). Fig 7 indicates that `RNAmountAlign` produces SPS scores comparable to `mLocARNA` and `LARA` and higher than `Multilign` and `FOLDALIGNM` while the SCI score obtained from `RNAmountAlign` are slightly lower than other software. Averaging over all sequences, the SPS scores for `RNAmountAlign, LARA, mLocARNA, FOLDALIGNM` and `Multilign` were respectively: 0.84 ± 0.17, 0.85 ± 0.17, 0.84 ± 0.17, 0.77 ± 0.22, and 0.84 ± 0.19. The left panel of Fig 8 indicates the run time of all software on a logarithmic scale, while the right panel shows the actual run time in seconds for `RNAmountAlign` as well as that of the next two fastest algorithms, `mLocARNA` and `LARA`. This figure clearly shows that `RNAmountAlign` has much faster run time than all other software in our benchmarking tests, thus confirming the earlier result from pairwise benchmarking.

**Fig 7.**
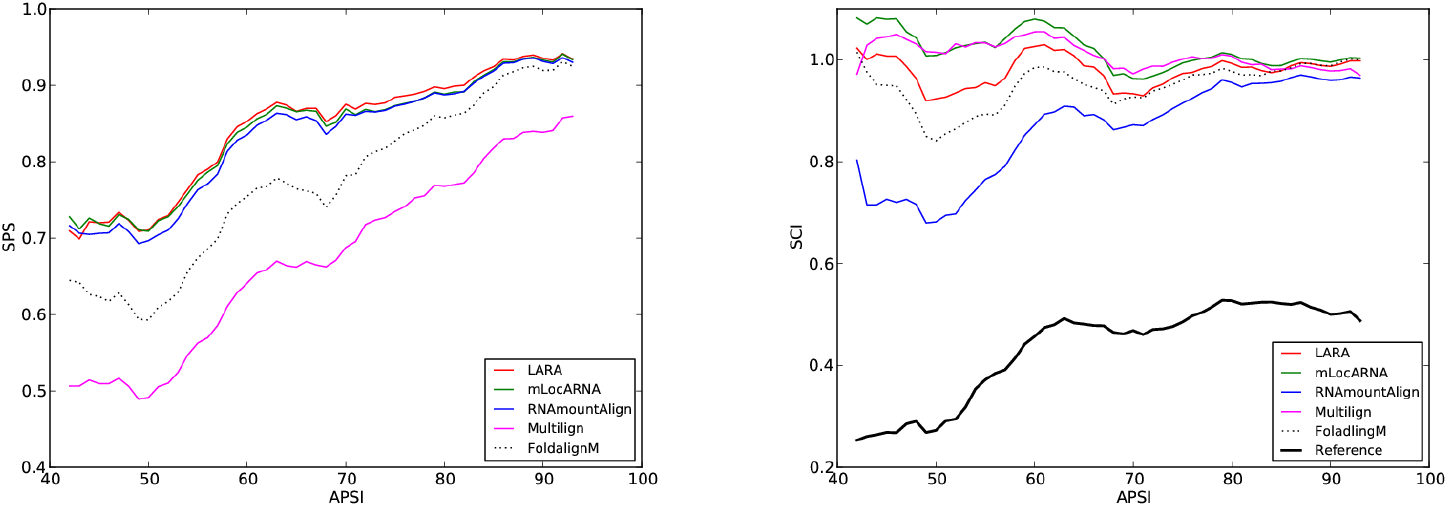
Sum-of-pairs(SPS) score (left) and structural conservation index (SCI) (right) for *multiple global alignments* using `RNAmountAlign, LARA, mLocARNA, FoldalignM` and `Multilign` . SPS and SCI are shown as a function of average pairwise sequence identity(APSI) in the k5 `BRAliBase 3` database used for benchmarking.

**Fig 8.**
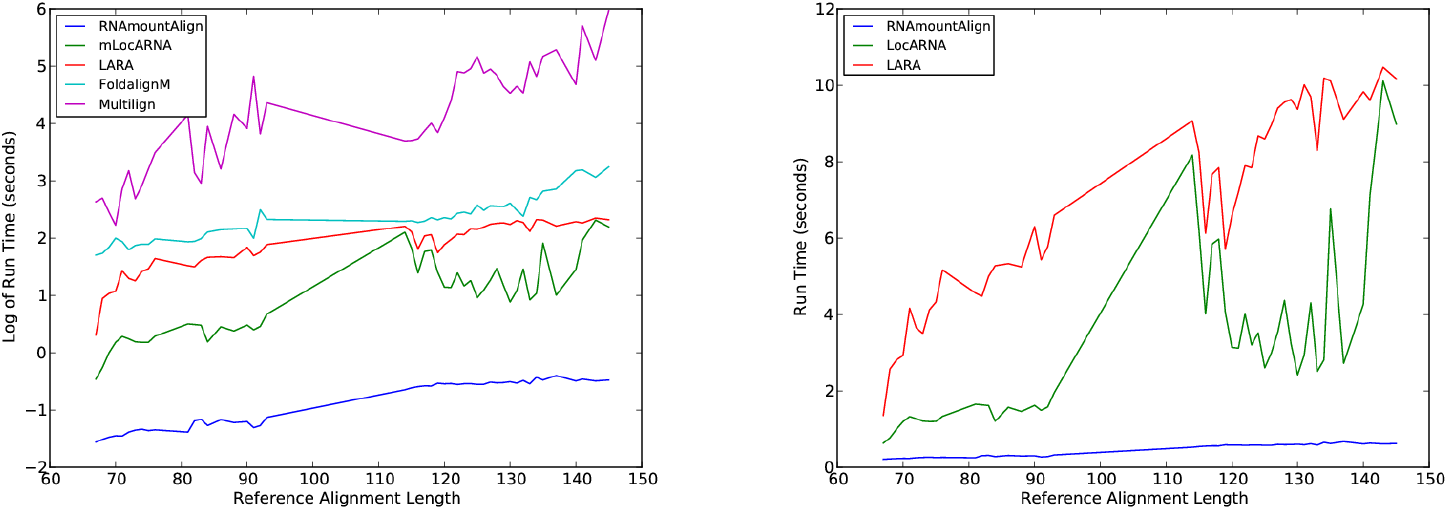
Run time of *multiple global alignment* for `RNAmountAlign, mLocARNA` and `LARA, FoldalignM` and `Multilign`. (Left) Log run time is as shown a function of reference alignment length for K5 alignments in `Bralibase 3.` (Right) Actual run time in seconds for `mLocARNA` and `LARA`.

## Conclusion

`RNAmountAlign` is a new `C++` software package for RNA local, global, and semiglobal sequence/structure alignment, which provides accuracy comparable with that of a number of widely used programs, but provides much faster run time. `RNAmountAlign` additionally computes E-values for local alignments, using Karlin-Altschul statistics, as well as *p*-values for normal, extreme value and gamma distributions by parameter fitting.

## Acknowledgements

Research supported by National Science Foundation grant DBI-1262439. Any opinions, findings, and conclusions or recommendations expressed in this material are those of the authors and do not necessarily reflect the views of the National Science Foundation.

## Footnotes

1 We follow [2,31] in our definition of mountain height, and related notions of ensemble mountain height and distance, while [32] and Vienna RNA package [4] differ in an inessential manner by defining *h_s_*(*k*) = |{(*i, j*) ∈ *s*: *i* < *k*}| – |{(*i, j*) ∈ *s*: *j* ≤ *k*}|.

2 Since we were unable to compile `STRAL`, our benchmarking results for `STRAL` use an adaptation of our code to support Eq (16). There are nevertheless some differences in how progressive alignment is implemented in `STRAL` that could affect run time.

